# Transdermal diagnosis and therapy using an integrated acoustofluidic patch

**DOI:** 10.64898/2026.09.09.750438

**Authors:** Yang Yang, Yantao Xing, Xiang Li, Zhuhao Wu, Hongwei Cai, Vivian Niu, Jack Crystal, Nian Wang, Ken Mackie, Weihua Guan, James Friend, Feng Guo

**Affiliations:** Department of Intelligent Systems Engineering, Indiana University, Bloomington, IN 47405, United States; Department of Medicine, Brigham and Women’s Hospital, Harvard Medical School, Boston, MA 02115, United States; Department of Engineering Physics and Department of Physics, Coe College, Cedar Rapids, IA, 52402, United States; Advanced Imaging Research Center, University of Texas Southwestern Medical Center, Dallas, TX 75390, United States; Gill Institute for Neuroscience, and Department of Psychological and Brain Sciences, Indiana University, Bloomington, IN 47405, United States; Department of Mechanical Engineering and Materials Science at Washington University in St. Louis, Louis, MO 63130, United States

**Keywords:** Personalized therapy, Acuate disease, Transdermal diagnosis, Transdermal therapy, Skin interstitial fluids, Acoustofluidics

## Abstract

Transdermal biosensing and drug delivery provide convenient, pain-free, and user-friendly solutions for personalized therapy. However, challenges remain in detecting and treating acute diseases, primarily due to current limitations in effective detection of transdermal biomarkers and administration of therapeutics to an individual within a short time window in a real-world environment. Here, we integrate a compact acoustofluidic patch for transdermal sampling, sensing, and delivery in a rapid and automated manner. By leveraging acoustic streaming and 3D-printed microfluidic designs, acoustofluidic sampler and injector modules are assembled for penetration of the skin stratum corneum, precise sampling of skin interstitial fluids, and controlled percutaneous release of therapeutics, respectively. By incorporating an electrochemical sensing module, the integrated patch can be further developed to achieve in situ and rapid detection of transdermal biomarkers and automated transdermal delivery of therapeutics via closed-loop control. As a proof-of-concept application, we demonstrate our approach can detect and reverse life-threatening acute allergic reactions in a mouse model of anaphylaxis by rapid sampling and sensing transdermal histamine followed by automated transdermal delivery of epinephrine. The innovative acoustofluidic patch may enable a new kind of transdermal devices for personalized therapy of various acute diseases.

---

Personalized therapy attracts tremendous attention in treating acute diseases because these conditions typically present with rapid onset, a short disease course, clearly defined symptoms, and a high risk of mortality. For instance, anaphylaxis is an unpredictable, acute, and potentially life-threatening allergic reaction affecting millions worldwide, especially children^1, 2^. Anaphylaxis occurs suddenly and may cause an individual to stop breathing, lose consciousness within minutes, and induce death in severe cases^1, 3^. Typically, within one hour of exposure to a specific allergen such as may occur in food, medications, etc^2^. In severe cases, an individual needs medical attention and therapy such as administration of epinephrine to prevent and relieve both airway obstruction caused by mucosal edema and/or bronchospasm, and hypotensive shock^4, 5^. However, it is challenging to accurately detect and rapidly manage such unpredictable complex systemic allergic reactions^7,8^. Currently, personalized therapy is still largely lacking for treating acute disease conditions within a specific time window, especially in real-world settings.

Transdermal devices hold promising potential for advancing personalized therapy against various diseases because theyprovide a minimally invasive interface to detect and manage the internal physiological environment of the human body^6-10^. So far, a wide range of transdermal drug delivery approaches have been developed and clinically deployed, particularly for treating chronic diseases^11-13^. Most clinically approved transdermal drug delivery systems rely on passive diffusion across the stratum corneum, which supports sustained release but lacks the temporal precision required for on-demand treatment in acute conditions^6, 14^. Moreover, a variety of physical enhancement strategies, such as iontophoresis^8^, microneedle-based patches^15^, acoustic-assisted delivery^11^, electroporation^16, 17^, and optically or thermally assisted approaches^18^, have been developed mainly for improving delivery efficiency, rather than for rapid and automated delivery of therapeutics. Thus, there is a critical need to innovate transdermal devices that enable rapid and automated drug delivery, with therapeutic decisions guided by accurate and timely diagnosis of acute disease conditions.

Transdermal biosensing may more accurately reflect systemic and tissue-level physiology than physiological signal (e.g., heart rate) sensing or sweat- and saliva-based measurements^19-21^. Transdermal biosensing typically accesses interstitial fluid (ISF) that represents a rich but underutilized source of diagnostic information^22^. Its molecular composition largely mirrors that of blood plasma while capturing localized and time-sensitive biochemical signals^23^. Moreover, compared with blood, biomarkers in ISF often exhibit faster changes and higher levels, highlighting its value for early detection of immune activation in acute pathological states^24-26^. A broad range of ISF biomarkers can be detected using emerging biosensing modalities, such as electrochemical enzymatic sensors for small molecules (e.g., glucose and lactate) ^27^ and electrochemical aptamer-based sensors for proteins and larger biomolecules^28, 29^. However, challenges remain for rapid, accurate, and in situ sensing of ISF biomarkers due to several practical limitations, including inefficient ISF extraction, complex sample handling, and reliance on bulky analytical instrumentation, all of which hinder their deployment to guide therapeutic decisions in real-world settings.

Acoustofluidics^30-38^, employing acoustics and microfluidics for handling biological samples and liquids, may address these unmet needs to innovate a new kind of active transdermal devices. Especially, our recent advances on digital acoustofluidic streaming^11, 39-41^ may provide a unique solution to extract and transport transdermal biomarkers for rapid and accurate transdermal biosensing, as well as a method to actively release therapeutics for rapid and automated transdermal drug delivery with digital regulation. Here, we innovate an integrated acoustofluidic patch for personalized therapy of acute disease by leveraging bidirectional acoustic streaming flows to facilitate minimally invasive transdermal diagnosis and therapy. We integrate sampling of transdermal fluids, transdermal biosensing, and transdermal drug delivery into a highly compact and portable patch device that (1) employs 3D-printed hollow microneedles to penetrate the skin barrier and establish transdermal microchannels for accessing subcutaneous fluid, (2) utilizes angle-controlled acoustic streaming to enable bidirectional fluid transport, allowing controlled ISF sampling and on-demand transdermal drug injection, (3) incorporates integrated biosensors for in situ analysis of the extracted samples, enabling real-time biomolecular diagnostics, and (4) implements a closed-loop management system to facilitate timely diagnosis and therapeutic intervention. Using an mouse model of anaphylaxis, we demonstrate that this acoustofluidic patch enables rapid assessment and timely therapeutic intervention, effectively reversing anaphylaxis without the need for manual drug administration. This integrated platform establishes a versatile framework for portable, personalized, and responsive management of acute diseases, highlighting the potential of acoustofluidic technologies for next-generation portable theranostics.

## Results

### Transdermal diagnosis and therapy using an integrated acoustofluidic patch

Here, we integrate a compact acoustofluidic patch implemented with a closed-loop control strategy to innovate transdermal devices for personalized therapy of anaphylaxis (**Fig. 1a**). This integrated device consists of two functionally distinct but electronically coordinated modules: (i) an acoustofluidic sampler with an in situ electrochemical sensor for rapid detection of anaphylaxis by actively sampling ISF and in situ sensing of ISF histamine, and (ii) an acoustofluidic injector for automated reversal of anaphylaxis through digital transdermal delivery of epinephrine with a closed-loop control (**Fig. 1b**). Specifically, the operational workflow is as follows: upon application to the skin, the hollow needle tips of the acoustofluidic sampler and injector can penetrate the stratum corneum, establishing microscale conduits between the microchannel and the dermal interstitial space. The sampler device uses acoustic streaming for jetting flow to actively draw ISF from the tissue into the embedded microchannel and sensing area. Meanwhile, the sampler device leverages acoustic streaming to induce rapid mixing and transport ISF efficiently toward the sensing electrodes of the integrated organic electrochemical transistor (OECT) sensor within the microchannel. When the detected concentration of biomarker (e.g., histamine) exceeds a predefined threshold, the system triggers the acoustofluidic injector via an implemented closed-loop control strategy. The injector device also uses acoustic streaming to drive therapeutic agents into the skin. In addition to directional programmable delivery, streaming vortices generated near the outlet promote local dispersion, facilitating drug distribution within the surrounding tissue. Both patches are applied to the skin and are governed by a shared control circuit, enabling coordinated operation in a closed-loop configuration (**Supplementary Fig. 1**).

**Fig. 1.**
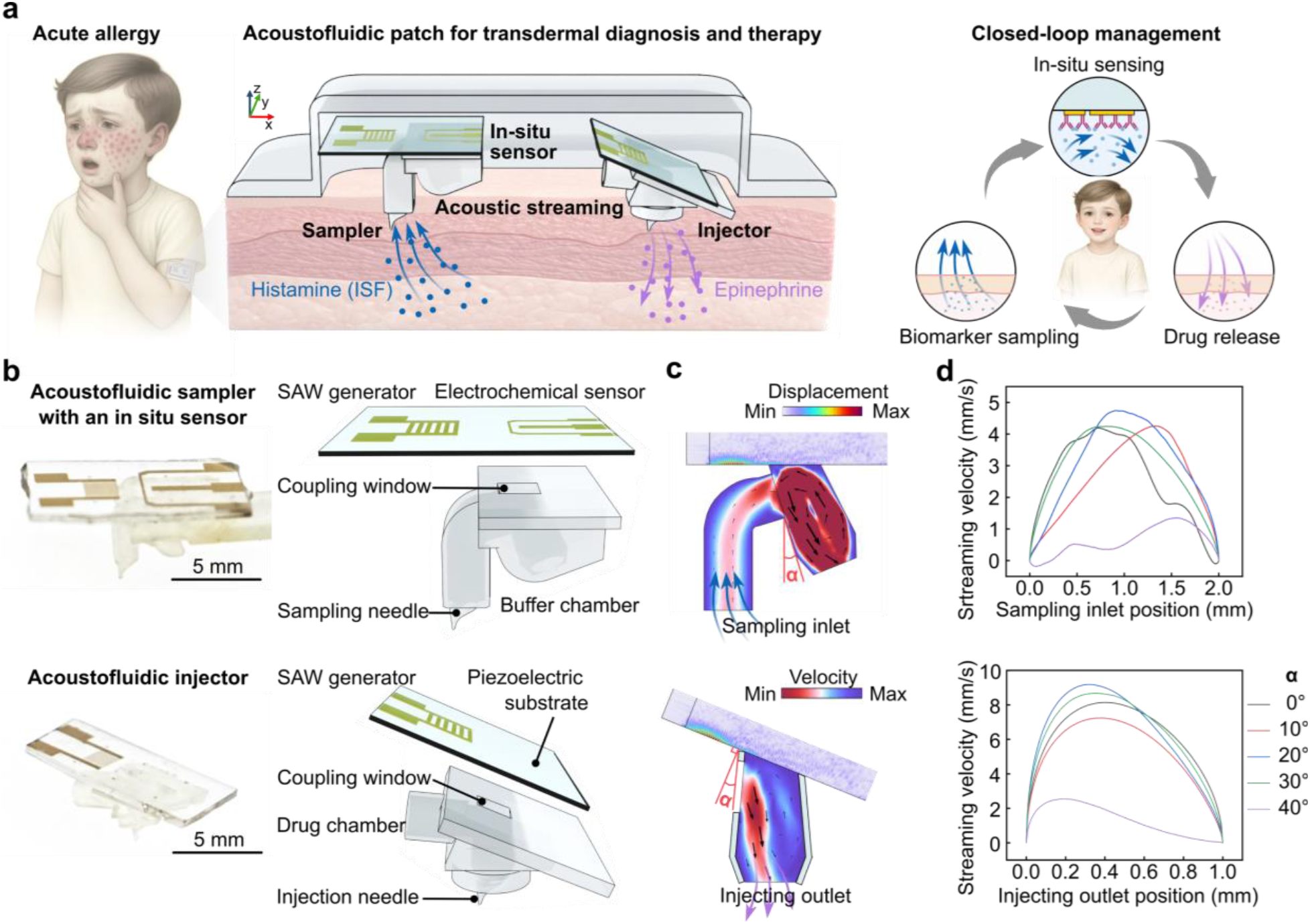
Transdermal diagnosis and therapy using an integrated acoustofluidic patch. (**a**) Schematics showing the working principle of an integrated acoustofluidic patch implemented with a closed-loop control strategy for personalized therapy of acute diseases (e.g., anaphylaxis). By leveraging digital acoustic streaming, an acoustofluidic sampler is designed to sample skin intestinal fluids (ISF) for in situ detection of transdermal biomarkers (e.g., histamine), and an acoustofluidic injector is developed to transdermally deliver therapeutics (e.g., epinephrine) for automated treatment of anaphylaxis. (**b**) Images and design diagrams of an acoustofluidic sampler device with an in situ electrochemical sensor and an acoustofluidic injector device. (**c**) Simulation results showing the displacement on the piezoelectric substrates and streaming within microchannels induced by surface acoustic waves (SAWs). (**d**) Simulation results of streaming velocity across the sampling inlet and the injecting outlet over different incident angles.

We employed acoustic streaming ^42, 43^, a kind of fluidic motion induced by sound waves, as an active pumping source both for sampling transdermal fluids and for releasing transdermal therapeutics. Stimulations were conducted to optimize the design of the acoustofluidic sampler and injector enabled by surface acoustic waves (SAWs). By deliberately engineering the incident angle between the propagating SAWs and the 3D-printed microchannels, we direct the resulting streaming jets either toward or away from the skin interface (micro-needles). This angle-controlled fluid flow enables single acoustofluidic architecture to operate in two distinct modes: inward-directed streaming for ISF extraction and outward-directed jetting for transdermal drug injection (**Fig. 1c**). Finite-element simulations were employed to visualize acoustic wave propagation and the resulting acoustic streaming fields, providing quantitative guidance for device design (**Supplementary Fig. 2a**). By systematically tuning the incident angle between the surface acoustic waves and the microchannel, we evaluated the dependence of streaming strength and directionality on acoustic coupling geometry (**Fig. 1d**). The simulations reveal that acoustic streaming is maximized when the incident angle approaches the Rayleigh angle (∼23 ° from a normal to the SAW generator’s surface), at which efficient mode conversion leads to strong momentum transfer into the fluid and pronounced jetting flows (**Supplementary Fig. 2b**). These results informed the geometric design of the microchannels to maximize acoustofluidic transport efficiency. With the optimized design, we further fabricated and tested the acoustofluidic sampler and injector for transdermal diagnosis and therapy.

### Active sampling of transdermal fluids using acoustofluidic streaming

The acoustofluidic sampler device was designed to actively extract interstitial fluid (ISF) carrying target biomarkers (e.g., histamine) into the sensing chamber using digitally-regulated acoustic streaming (**Fig. 2a**). Here, we fabricated and characterized the acoustofluidic sampler device in vitro. To visualize the microscale acoustic streaming, fluorescent tracer particles were introduced into the sampling microchannel (**Fig. 2b**). Time-stacked particle trajectory images clearly reveal inward-directed streamlines at the microneedle inlet, confirming active fluid extraction into the sensing chamber with acoustic streaming (Streaming (+)). Within the chamber, vortical flow patterns were observed, indicating rapid convective mixing induced by acoustic streaming. In contrast, when acoustic streaming is off (Streaming (-)), tracer particles remained stationary, demonstrating the negligible leakage under passive conditions. A linear relationship was observed between the sampling flow rate and the input voltage (**Fig. 2c**), providing a quantitative basis for precise and predictable control of the sampling process. Then, we characterized post-penetration skin recovery in mice, confirming minimal, reversible tissue perturbation (**Supplementary Fig. 3**). We further assessed the thermal response of the device. By balancing performance with thermal safety, a pulsed operation mode (50% duty cycle, 17 V_pp_) was selected as the optimal condition for subsequent experiments, maintaining the operating temperature below physiological body temperature to avoid thermal injury (**Supplementary Fig. 4**)^44^.

**Fig. 2.**
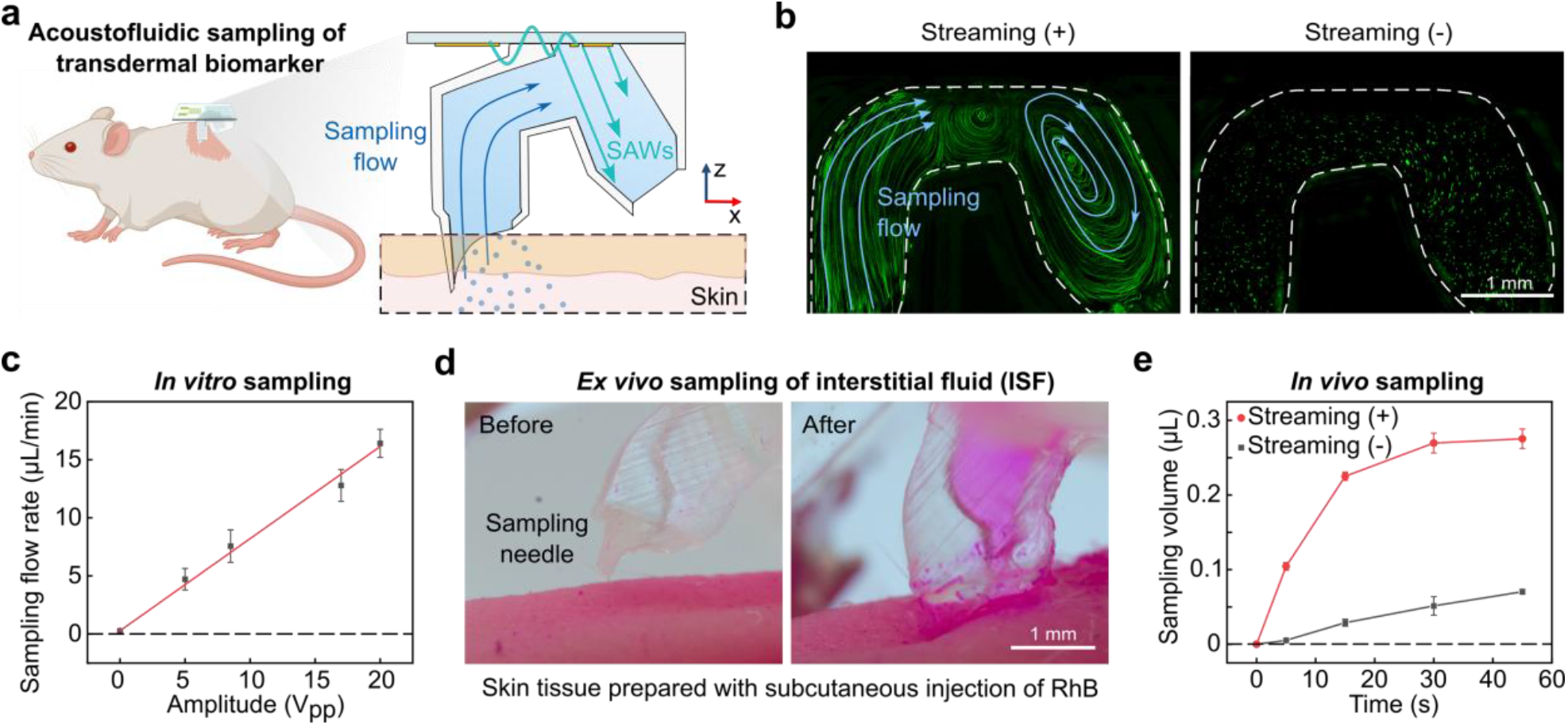
Active sampling of transdermal fluids using acoustofluidic streaming. (**a**) Schematic illustration showing acoustofluidic sampling of transdermal biomarkers in vivo. (**b**) Stacked fluorescent particle trajectories indicating flow profiles within the chamber of an acoustofluidic sampler with (streaming (+)) or without (streaming (-)) streaming. Scale bar: 1 mm. (**c**) *In vitro* characterization of the sampling flow rate over input acoustic amplitude (V_pp_). Data are presented as mean ± SD; n = 5 independent experiments. (**d**) Images before and during the acoustofluidic sampling of transdermal ISF (pre-subcutaneous injection of Rhodamine B for visualization) from a mouse tissue (*ex vivo* model). Scale bar: 1 mm. (**e**) *In vivo* quantification of accumulated sampling volume over time with (streaming (+)) or without (streaming (-)) streaming. Data are presented as mean ± SD; n = 4 independent experiments.

Next, we tested transdermal fluid sampling on *ex vivo* biological tissue. The acoustofluidic sampler was applied to pretreated mouse skin tissues with subcutaneous injection of Rhodamine B solution for visualization. With acoustic streaming, a rapid influx of dye-labeled interstitial fluid into the patch microchannel was observed, indicating efficient acoustofluidic extraction during sampling (**Fig. 2d, Supplementary Fig. 6a**,**b**). Furthermore, we quantitatively evaluated the sampling efficiency *in vivo* using a mouse model. The extracted ISF volume was calculated by measuring the total amount of extracted glucose in the acoustofluidic sampler and comparing it with the simultaneously measured concentration of blood glucose in the same mouse. Turning on the acoustic actuation led to a clear increase in sampling efficiency compared with the streaming OFF condition (**Fig. 2e**). Notably, a sufficient volume of ISF for biomarker detection was collected within 1 min, satisfying the timeliness requirements for rapid assessment in acute settings. The *in vivo* extraction profile differs substantially from that observed under *in vitro* conditions. First, the higher hydraulic resistance imposed by skin and subcutaneous tissues results in a markedly reduced extraction rate. Second, the *in vivo* extraction curve deviates from linearity, in contrast to the approximately linear trend observed *in vitro*. This nonlinearity is likely attributable to localized depletion of ISF in the vicinity of the sampling interface during continuous extraction. Collectively, these results demonstrate that the acoustofluidic sampler enables rapid, controllable, and minimally invasive ISF sampling, providing a robust foundation for subsequent *in situ* biomarker sensing.

### In-situ electrochemical sensing of transdermal biomarker

Following the acoustofluidic sampling and transportation of ISF biomarkers to the sensing chamber, we next characterized the OECT biosensor integrated within the acoustofluidic sampler for in situ detection of anaphylaxis. Since histamine is a ubiquitous mediator for allergic reactions^45^, we chose ISF histamine for in situ detection of anaphylaxis. However, despite its high specificity, histamine is rarely used as a routine clinical diagnostic marker, because current clinical diagnostic methods and systems have difficulty in effectively measuring it during its rapid rise and fall following allergen exposure^46^. To address the challenge, an in situ OECT sensor functionalized with histamine antibodies was embedded directly within the sampling chamber, allowing direct exposure and rapid measurement of freshly extracted ISF (**Fig. 3a**). The trajectories of fluorescent tracers further indicate that acoustofluidic streaming not only drives ISF extraction but also enhances sensing by convectively transporting target molecules toward the sensor interface (**Fig. 3b**). By using histamine standards at varying concentrations, we characterized the response of the OECT sensor with/without antibody (**Fig. 3c**). Histamine binding to the functionalized sensor induced systematic shifts in the transfer characteristics, whereas non-functionalized devices exhibited negligible responses. Quantitative analysis of the normalized current change (ΔI_ds_/I_ds(0)_) revealed a concentration-dependent response over a clinically relevant range of histamine concentrations, enabling construction of a calibration curve (**Fig. 3d**)^47, 48^. As a control, the device without antibody functionalization exhibited no detectable response across the tested histamine concentrations. These results show that the integrated sensor enables reliable *in situ* histamine detection over a broad concentration range, with sufficient sensitivity and selectivity for threshold-based analysis.

**Fig. 3.**
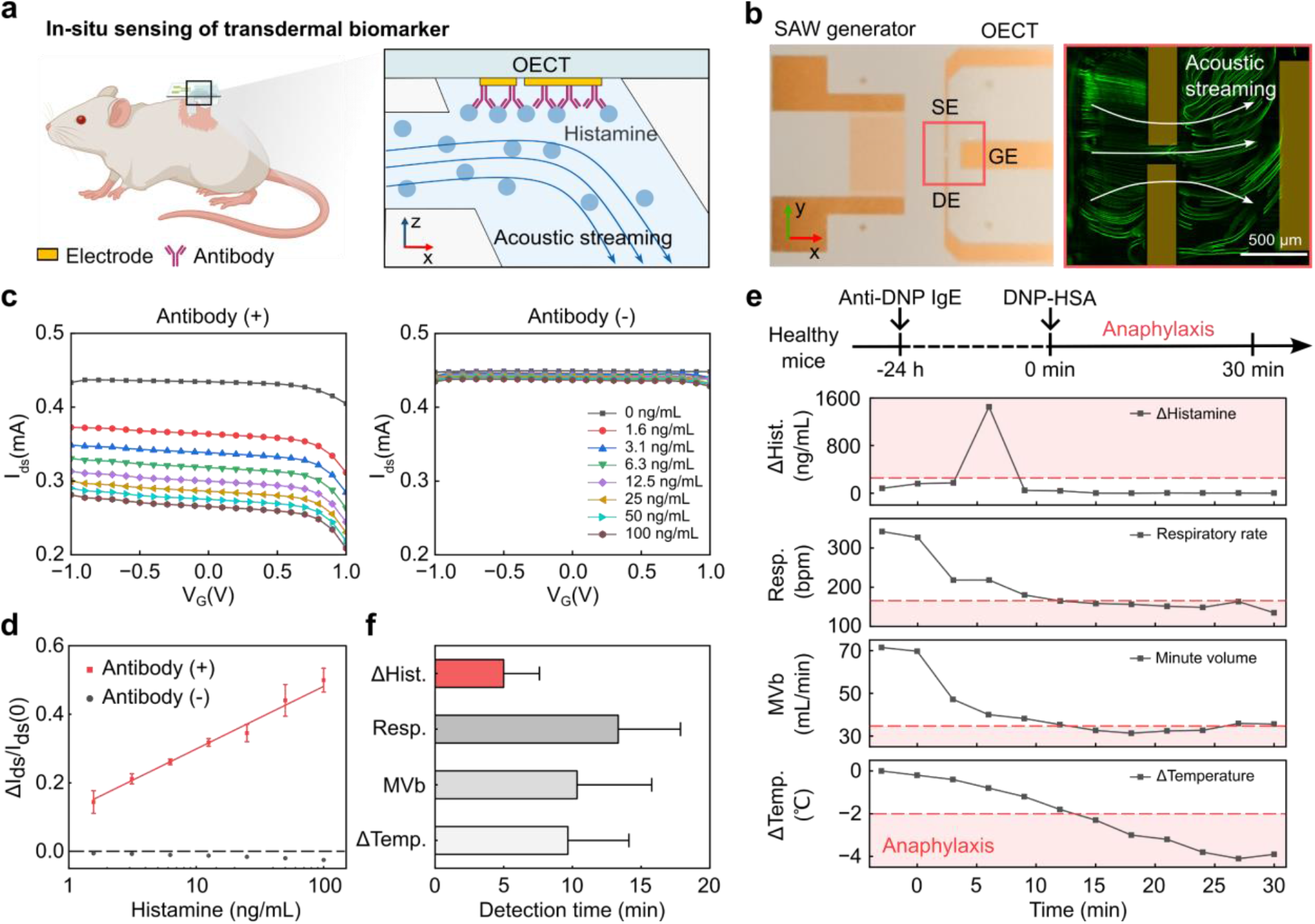
In-situ electrochemical sensing of transdermal biomarkers. (**a**) Schematic diagram showing in-situ sensing of biomarkers (e.g., histamine) using an integrated organic electrochemical transistor (OECT). Acoustic streaming transports histamine toward antibody-functionalized electrodes for real-time detection. (**b**) Bright-field and fluorescence flow visualization images showing the efficient transport of the sample across the detection electrode using acoustic streaming. (**c**) Transfer characteristics of the OECT under increasing histamine concentration, showing a concentration-dependent shift in drain current (I_ds_). (**d**) Histamine dose-response curve demonstrating selective detection with antibody-modified sensors compared to a negligible response without antibody. Data are presented as mean ± SD; n = 5 independent experiments. (**e**) Schematic of IgE-dependent anaphylaxis induction in mice. *In vivo* measurement of histamine levels, respiratory rate, minute volume (MVb), and body temperature change in a mouse model of anaphylaxis. Histamine rises rapidly post-challenge, reaching the anaphylactic threshold before physiological parameters. (**f**) Comparison of detection time for histamine, temperature drop, and respiratory distress, indicating earlier detection capability of the histamine sensor (<10 min). Data are presented as mean ± SD; n = 9 independent mice.

We next cascaded acoustofluidic sampling with in-situ sensing to evaluate the performance of the acoustofluidic sampler in a mouse model of anaphylaxis. A mouse model of IgE-mediated anaphylaxis was established by sensitization with anti-DNP IgE, followed by systemic challenge with DNP–HSA to induce rapid mast cell degranulation and histamine release (**Fig. 3e**)^49-51^. We measured histamine levels in ISF and serum at multiple time points in the anaphylaxis mouse model, providing further validation that ISF sampling is sufficient to reflect systemic histamine dynamics (**Supplementary Fig. 6**). The acoustofluidic sampler was applied to mice to continuously monitor histamine levels in ISF during the onset of anaphylaxis. Strikingly, the acoustofluidic sampler detected a sharp rise in histamine concentration shortly after exposure, well before severe physiological symptoms had fully developed (**Fig. 3e, Supplementary Fig. 7**). In parallel, respiratory depression and reduced body temperature were quantified in the same anaphylaxis model. Comparison of these readouts highlights the temporal advantage of histamine sensing over physiological indicators, while also underscoring its higher biochemical specificity (**Fig. 3f**). These results show that combining acoustofluidic sampling with integrated OECT sensing enables time-resolved *in situ* histamine monitoring *in vivo* (<10 min), allowing early biochemical changes that are often missed by conventional clinical measurements to be captured and used to guide therapy.

### Closed-loop reversal of anaphylaxis via acoustofluidic delivery of epinephrine

We next integrated a closed-loop system with our well-characterized acoustofluidic sampling, in situ sensing, and delivery modules (**Fig. 4a**). The patch implemented with the closed-loop control is designed to detect and treat allergic reactions enabling automated and rapid reversal of anaphylaxis. Acoustofluidic sampling of ISF, in situ sensing of ISF histamine, and acoustofluidic injection of epinephrine operate in a cascaded workflow. Once the detected histamine level exceeds a predefined threshold, the injector is triggered to deliver epinephrine for anaphylaxis treatment. Before validating the closed-loop system for automated reversal of anaphylaxis, we characterized the acoustofluidic delivery in vitro. We similarly employed a fluorescent tracer to visualize fluid transport during acoustofluidic injection (**Fig. 4b**). An outward-directed jetting was observed at the injector outlet under streaming-on conditions.

**Fig. 4.**
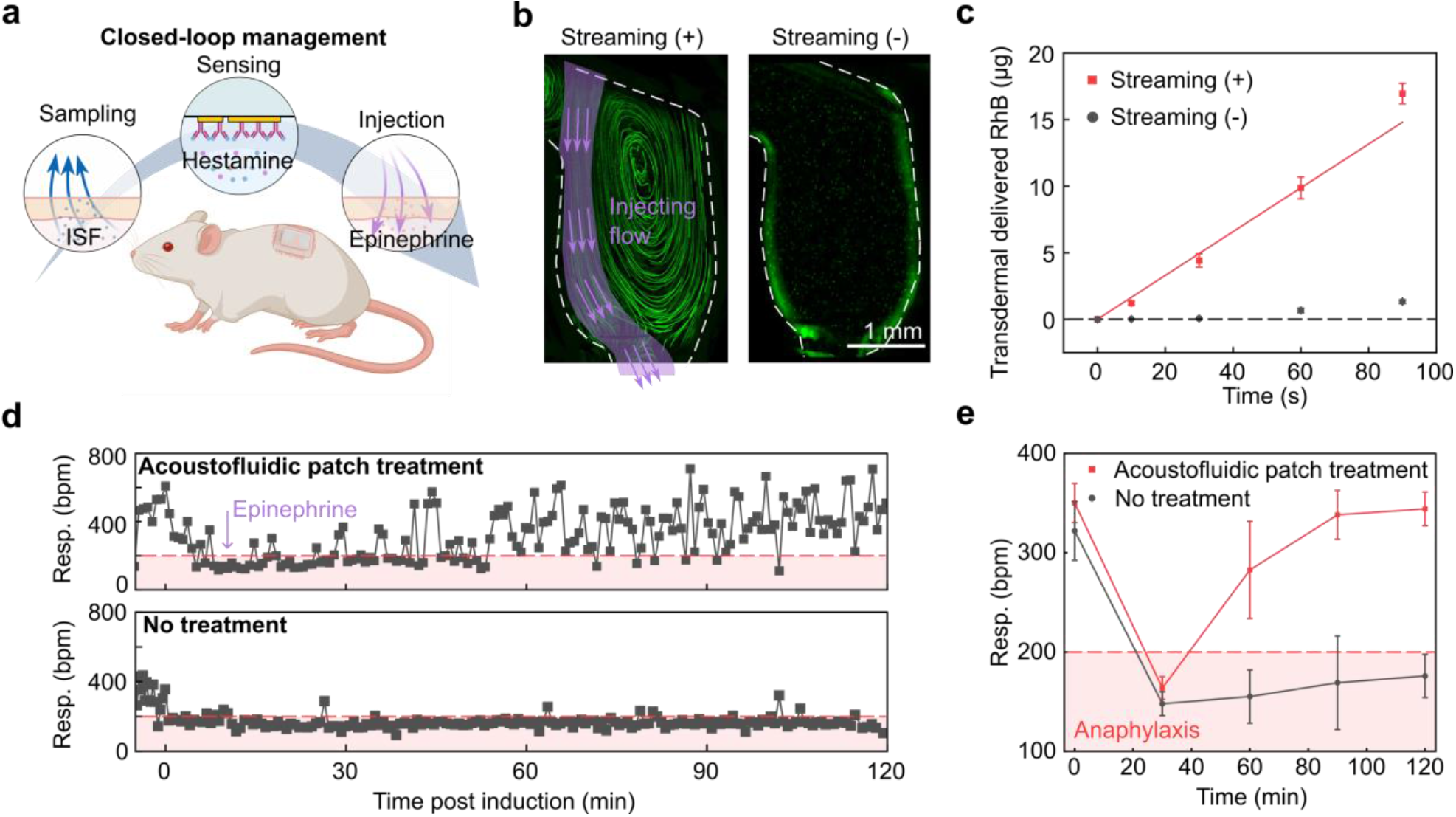
Closed-loop reversal of anaphylaxis via acoustofluidic delivery of epinephrine. (**a**) Schematic diagram showing the closed-loop management of anaphylaxis through an integrated acoustofluidic patch. (**b**) Visualization of jetting flows in the injection chamber with (streaming (+)) or without (streaming (-)) streaming. (**c**) Quantification of transdermally delivered Rhodamine B dye over time with (streaming (+)) or without (streaming (-)) streaming, indicating the significantly enhanced transdermal delivery mediated by active acoustic streaming-mediated release. Data are presented as mean ± SD; n = 5 independent experiments. (**d**) Reversal of anaphylaxis using acoustofluidic patch treatment in vivo. (**e**) Respiratory rate post-anaphylaxis with and without acoustofluidic patch treatment. Mice receiving an acoustofluidic patch treatment showed significant recovery compared to the control group. The shaded area indicates the healthy respiratory condition. Data are presented as mean ± SD; n = 9 independent mice.

In contrast to the sampling configuration, the design of the injector allows SAWs to propagate through the liquid medium toward the microneedle–dermal interface. Moreover, the ability of acoustic waves to penetrate biological tissues makes them suitable for intratissue delivery^52-55^. At this interface, acoustic excitation induces secondary vortical flows and localized nebulization effects (**Supplementary Video 1**). The acoustic losses in the polymer of the microneedles and in the dermal tissue ensure that the acoustic (Poynting) power flows from the SAW source toward the tissue via the needles, leading to acoustic streaming along these same directions. As the acoustic energy propagates further into the tissue, it may facilitate dispersion and uptake of the delivered payload within the dermal tissue.. We further quantified injection performance by acoustofluidic releasing Rhodamine dye into *ex vivo* mouse skin tissues over different treatment times. The quantification results showed the linear increase of transdermal delivered Rhodamine dye over time under active delivery via acoustic streaming (streaming (+)), while very little Rhodamine dye was transdermally delivered under passive delivery without acoustic streaming (streaming (-)) (**Fig. 4c**). Thus, our results demonstrate the successful transdermal delivery of histamine for reversal of anaphylaxis in vivo.

We further evaluated the efficacy of the closed-loop acoustofluidic patch treatment in a mouse model of anaphylaxis. In the treatment group, mice were equipped with the acoustofluidic patch and subsequently induced into anaphylaxis. Upon detection of a critical rise in ISF histamine, the patch system triggered acoustofluidic delivery of epinephrine through the injection patch. A control group of mice underwent the same anaphylaxis protocol without receiving treatment. The respiratory rate was continuously monitored as an indicator of the allergic state of the mice. Respiratory function recovered after acoustofluidic injection of epinephrine in the mice with acoustofluidic patch treatment, whereas untreated animals showed progressive respiratory depression (**Fig. 4d**). Group-level analysis (n=9) further confirmed that the acoustofluidic patch treatment significantly improved respiratory outcomes compared with no-treatment controls (**Fig. 4e** and **Supplementary Fig. 8**). Together, the results demonstrate the successful integration of acoustofluidic patch with a closed-loop control for automated and rapid reversal of anaphylaxis.

## Discussion

In this study, we developed a closed-loop theranostic patch that integrates acoustofluidic sampling, on-patch biosensing, and acoustofluidic injection within a single wearable platform. By engineering the spatial coupling between surface acoustic waves and three-dimensional microchannels, we established an angle-controlled acoustofluidic strategy that enables programmable, bidirectional transport across the skin barrier using a unified physical mechanism. By coupling early biochemical readouts to timely therapeutic delivery, this system enables feedback-controlled intervention, representing a transition from symptom-controlled management toward biomarker-triggered treatment in acute conditions.

Despite these advances, several challenges remain before translation beyond proof-of-concept demonstrations. Further optimization is needed to improve system-level integration, including long-term stability of therapeutic payloads, robustness of on-demand skin penetration, and reliability under repeated operation in real-world environments. In addition, practical deployment will require intuitive user interfaces, enhanced material durability, and integrated wireless control to ensure safe and effective use outside controlled laboratory settings. Addressing these challenges will be critical for advancing acoustofluidic theranostic patches toward clinical and consumer-facing applications. Moreover, a key barrier to use of the technology is FDA approval of these devices, which may be uniquely challenging due to the introduction of physiologic closed-loop control: an automated diagnostic function that delivers medication in response to a sensed condition in an integrated device for a condition that—in most cases—has traditionally required clinical emergent evaluation^56^.

Overall, this study establishes acoustofluidics as a versatile framework for real-time transdermal theranostics. By enabling a closed-loop strategy, the presented patch provides a foundation for patient-centered platforms capable of timely point-of-care diagnosis and treatment, particularly for acute medical conditions. More broadly, acoustofluidic transdermal technology occupies a unique position between highly invasive approaches, such as deeply penetrating needles or device implantation, and noninvasive modalities based on sweat analysis or electrophysiological monitoring. This intermediate positioning offers a favorable balance between invasiveness and performance, enabling minimally invasive operation suitable for short-term, repeated operation while maintaining high sensitivity and specificity through direct analysis of molecular targets in the ISF and highly efficient drug delivery. Thus, such a platform may be further extended to the management of chronic conditions, including glucose monitoring and pain management, highlighting the broader potential of acoustofluidic transdermal systems for long-term, adaptive healthcare applications.

## Methods

### Simulation of acoustofluidic streaming

Acoustic streaming induced by attenuated surface SAWs was simulated using COMSOL Multiphysics 6.3 using our early developed pipelines^57-59^. The piezoelectric LiNbO_3_ substrate was modeled using coupled electrostatics and solid mechanics modules to capture SAW excitation under an applied alternating voltage. Wave propagation within the fluid domain was resolved using the pressure acoustics interface in the frequency domain, while steady streaming velocities arising from acoustic attenuation were computed using the creeping flow interface. Multiphysics coupling between the acoustic and flow fields was implemented through the acoustic streaming domain coupling module. Perfectly matched layers were applied at the substrate boundaries to suppress spurious wave reflections. This simulation framework enabled quantitative evaluation of streaming flow patterns as a function of SAW incidence angle and driving voltage (**Fig.1** and **Supplementary Fig. 1**).

### 3D printing of acoustofluidic hollow needle and streaming chamber

Acoustic streaming chamber and hollow microneedles were fabricated using a commercial stereolithography (SLA) 3D printer (Sonic Mini 8K, Phrozen, Hsinchu City, Taiwan, China). The original 3D designs were created using Autodesk Fusion 360 (version 2.0.12670, Autodesk Inc, USA) and subsequently exported to slicing software (Chitubox, version 1.9.4, CBD-Tech, China). Following printing, the components underwent a 10-minute washing step in isopropyl alcohol (IPA, 67-63-0, EMD Millipore, Burlington, USA). Subsequently, the devices were air-dried and further curing using 405 nm light at 60°C for 30 minutes to promote additional crosslinking and enhance the mechanical strength of the microneedles.

### Fabrication of SAW generators and OECTs

SAW generators and OECTs were fabricated on a 128° Y-cut lithium niobate (LiNbO_3_) wafer by depositing a Cr/Au layer (5 nm/200 nm) using thermal-evaporation (Auto 306, Boc Edwards, USA), following our previously well-developed procedures^39, 60, 61^. Interdigital transducers (IDTs) of SAW generators consisted of 15 electrode finger pairs with a finger width and gap of 25 µm with a consequent resonant frequency of 37.4 MHz, yielding an aperture of 2.5 mm. For the OECT architecture, the organic channel was designed with a length and width of 160 µm × 100 µm. The lateral gate electrode (1 mm in width) was positioned 500 µm away from the channel. Electrode patterns for both IDTs and OECTs were defined by standard photolithography using a chrome film mask (Artnet Pro, Inc., USA) and photoresist AZ5214E (Kayaku Advanced Materials, Inc., USA). After metal deposition, excess metal was removed via a lift-off process using Remover PG (Kayaku Advanced Materials, Inc., USA), followed by rinsing with acetone. Electrical connections were established by bonding external wires to the electrodes using silver epoxy (MG Chemicals, USA).

### Functionalization of OECTs

Following oxygen plasma cleaning (PDC-001, Harrick Plasma, USA), a thin film of poly(3,4-ethylenedioxythiophene):poly(styrenesulfonic acid) (PEDOT:PSS) was spin-coated onto the organic channel at 2000 rpm for 60 s, yielding a film thickness of approximately 100 nm. The coated devices were subsequently annealed at 80 °C for 1 h under a constant nitrogen purge. Histamine polyclonal antibodies (ENZ-ABS292-0100, Enzo Life Sciences, Inc., Farmingdale, USA) were immobilized onto the PEDOT:PSS surface via silanization using a 2% glycidoxypropyltrimethoxysilane (GPMS) solution in toluene, followed by overnight incubation at 4 °C. After functionalization, the devices were rinsed three times with 1× phosphate-buffered saline (PBS) and blocked with 2% bovine serum albumin (BSA) for at least 30 min to minimize nonspecific binding. This fabrication protocol was adapted from previously reported methods^62, 63^. Laser-cut transfer tapes were used as masks to define the regions for organic layer deposition and antibody functionalization.

### Assembly and operation of the acoustofluidic patch

The acoustic streaming chamber was bonded directly onto the LiNbO_3_ substrate using a 130 µm–thick transfer tape (468MP, 3M, USA) as the adhesive layer. To minimize acoustic energy loss due to dissipation, the bonding layer was patterned into a rectangular geometry with a thin wall (0.75 mm) oriented toward the propagation direction of SAWs. The chambers were pre-filled with 1× phosphate-buffered saline (PBS) prior to operation. For acoustofluidic injection, polytetrafluoroethylene tubing (Cole-Parmer, USA) was connected to the chamber inlet as a drug reservoir. The acoustofluidic patch devices were actuated using a sinusoidal radiofrequency signal (Freq= 37.4 MHz, V_pp_ = 17 V, 50% duty cycle) generated by a function generator (TGP3152, Aim TTi, UK) and amplified with a power amplifier (LZY-22+, Mini-Circuits, USA).

### Characterization and calibration of OECTs

The transfer characteristics of the OECT devices were measured using source-measure units (Keithley 2612B, Tektronix, USA). Before and after each in situ histamine detection step, the devices were thoroughly rinsed three times with 1× PBS. Electrical measurements were performed at a constant source–drain voltage (V_ds_ = −0.1 V) while sweeping the gate voltage (V_G_) from −1 to 1 V. The relative change in source–drain current (ΔI_ds_/I_ds0_) was calculated following the antigen–antibody binding reaction. Each transfer curve measurement was repeated five times to ensure reproducibility. Histamine-spiked saline solutions were employed as calibration standards to quantify the relationship between sensor readout and histamine concentration.

### Electronic system design and integration

A two-layer flexible printed circuit board (FPCB) was designed using Altium Designer, with a rounded rectangular outline (31 mm × 40 mm) matching the footprint of the acoustofluidic sensor patch. The electronic system integrates voltage regulators (TPS60403, Texas Instruments; XC6206P33, Torex Semiconductor) for power management, a boost converter (LM2735, Texas Instruments), an electrochemical front end (AD5941, Analog Devices), a 37.4 MHz crystal oscillator for excitation signal generation, a MOSFET (IRFZ48N) for signal amplification, and a Bluetooth Low Energy (BLE) module (RF-BM-BG22C3, RF-STAR) for system control and wireless communication. Power was supplied by a rechargeable 3.7 V lithium battery with a capacity of 40 mAh.

### Characterization of acoustofluidic streaming

To visualize acoustofluidic streaming patterns, the acoustic chamber was designed with a sidewall opening and sealed with a coverslip. Fluorescent polystyrene beads (FSDG005, Bangs Laboratories Inc., USA) were used as tracer particles, and videos were recorded using an inverted fluorescence microscope (IX-81, Olympus, Japan). Tracer streamlines were reconstructed from maximum-intensity projections of every 15 frames using the open-source ImageJ plugin Flowtrace^64^. To quantify the volumetric flow rate induced by acoustic streaming, an auxiliary reservoir was incorporated around the hollow microneedle tip and pre-filled with 100 µL of deionized (DI) water. To eliminate gravitational or siphoning effects, the tubing inlet and needle tip were positioned at the same height. By varying the duration of acoustic actuation, the transported fluid volume was determined by multiplying the measured displacement of the fluid along the tubing by its cross-sectional area. When acoustic actuation was inactive, convective flow was negligible; therefore, needle patches were pre-filled with a colored dye, and diffusion-driven color changes in the reservoir were monitored over time. The equivalent diffused volume was subsequently quantified using a calibration curve.

### Closed-loop control strategy

A closed-loop acoustofluidic system was developed to integrate sampling, biochemical sensing, and therapeutic delivery within a wearable patch. In the sampling module, a radio-frequency excitation circuit drives a SAW transducer to generate localized acoustic streaming, actively enhancing ISF transport toward the sensing interface. Histamine concentration is monitored using an organic electrochemical transistor (OECT)-based sensor, where analyte-induced ionic modulation of the channel conductance is transduced into electrical signals and digitized by an integrated electrochemical front end for real-time analysis. When the measured histamine signal exceeds a predefined threshold, the control unit autonomously triggers a secondary acoustofluidic actuation module (injecting module). This module activates a dedicated SAW generator coupled to the acoustofluidic injector, enabling rapid and controlled transdermal release of epinephrine. The system operates as a sampling-sensing-decision-making-action loop, allowing autonomous biochemical detection and immediate therapeutic intervention without external supervision, thereby establishing a fully integrated closed -loop portable platform.

### Animal models of anaphylaxis

All animal procedures were performed in accordance with the National Institutes of Health Guidelines for the Care and Use of Laboratory Animals and were approved by the Institutional Animal Care and Use Committee (IACUC) at Indiana University Bloomington (Protocol NO. 22-006). All efforts were made to minimize animal suffering and reduce the number of animals used. BALB/c mice (20–30 g) were purchased from Envigo (Indianapolis, IN, USA) and housed in a controlled environment (temperature and humidity) with HEPA-filtered air under a 12 h light/dark cycle (lights on from 7:00 AM to 7:00 PM), with food and water provided ad libitum. A passive systemic anaphylaxis (PSA) model was established as previously reported^49-51^. Briefly, mice were sensitized via intraperitoneal injection of 200 µL saline containing 100 µg/mL monoclonal anti-dinitrophenyl (anti-DNP) IgE antibody. After 24 h, anaphylaxis was induced by intravenous injection of 100 µL saline containing 10 mg/mL dinitrophenyl–human serum albumin (DNP–HSA). Throughout the anaphylaxis induction and treatment period, respiratory rate was continuously monitored using whole-body plethysmography (WBP) (Buxco Small Animal WBP, Data Sciences International, USA), while the temperature was monitored using a rectal temperature probe. Specifically, a histamine concentration exceeding 200 ng mL^−1^ was used as a threshold indicative of anaphylaxis. Respiratory depression was defined as a reduction in respiratory rate or baseline minute ventilation (MVb) to ≤50% of the individual baseline, reflecting clinically relevant ventilatory compromise. Acute hypothermia, a well-established hallmark of systemic anaphylaxis in murine models, was defined as a drop in core body temperature of at least 2 °C relative to baseline.

### Ex vivo and in vivo evaluation of acoustofluidic patches

For both *ex vivo* and *in vivo* experiments, fur on the dorsal skin was removed under isoflurane anesthesia (4% induction, 2 – 2.5% maintenance), with body temperature maintained using a heating pad. Fur removal was performed by mechanical trimming followed by a brief application of depilatory cream, which was removed after ∼2 min by thoroughly washing with warm saline. For *ex vivo* sampling studies, mice were euthanized after fur removal. Freshly harvested skin tissues were used as an *ex vivo* experimental model. A total of 100 µL phosphate-buffered saline (PBS) containing Rhodamine B (25 µg/mL) was injected subcutaneously as a fluorescent tracer. The sampling patch was applied at a location more than 5 mm away from the injection site. Sampling efficiency was evaluated by fluorescence intensity within microchannels using an inverted fluorescence microscope (IX-81, Olympus, Japan). For *ex vivo* injection studies, Rhodamine B (0.4 mg/mL) was used as a fluorescent tracer to visualize the delivery process. The total injected volume was quantified by multiplying the displacement of the liquid front within the injection tubing by the cross-sectional area of the tubing. For *in vivo* sampling and injection studies, mice were fully anesthetized, and the acoustofluidic patches were stably affixed to the exposed dorsal skin using a custom 3D-printed applicator. The extracted ISF was diluted and analyzed using a commercial assay kit (Invitrogen A22189, Thermo Fisher Scientific, USA). By comparing the total glucose amount measured in the extracted ISF with that in 1 µL of serum simultaneously collected from the tail vein, the effective volume of ISF extracted by the acoustofluidic sampler was calculated.

### Histamine detection and intervention during anaphylaxis

Following induction of anaphylaxis, ISF was sampled using the acoustofluidic patch at 5 min intervals, and in situ histamine detection was performed sequentially using pre-functionalized and integrated OECT biosensors. For each sensor, a four-point calibration curve was established to quantify histamine concentration at different time points. Histamine standards of 0 and 0.5 ng/mL were applied prior to ISF sampling, while standards of 100 and 200 ng/mL were applied after completion of the sequential sampling and detection process. Temporal concordance between histamine levels in interstitial fluid and serum was independently validated using a commercial histamine ELISA kit (IT6088, G-Biosciences, USA) following the manufacturer’s protocol (**Supplementary Fig. 6**). For therapeutic intervention, the acoustofluidic injector was preloaded with epinephrine (0.1 mg/mL) diluted in saline. Upon detection of an anaphylactic signal exceeding the predefined threshold, the closed-loop controller triggered the acoustofluidic injector to initiate a 2-min transdermal drug delivery, achieving a targeted delivery volume of 50 µL. Respiratory rate was continuously recorded until the baseline rate was restored.

## Data availability

Experimental data supporting the findings of this study are available from the corresponding author upon reasonable request. The dataset generated in this study, which is necessary to interpret, verify, and extend the article’s findings, is provided in the Supplementary Information/Source Data file. Source data are provided in this paper.

## Acknowledgments

F. G. acknowledges the National Institutes of Health grants (U01DA056242, U54AG090792, R01GM160423, and R01DK133864) and National Science Foundation (EFRI2422149). K.M. acknowledges the National Institutes of Health grants (U01DA056242 and R01AT061112)

## Author contributions

F.G., J. F., and W.G. conceived and designed the project; Y. Y. and X.L. fabricated and characterized the device with help from Z. W., J. X., V.N., and J.C.; Y.X. developed the closed-loop system; Y.X. and X.L. performed the animal experiments for device and system validation; Y. Y., Y. X., X.L., H. C., N. W., K.M., W.G., J. F., and F.G. analyzed the data; Y.Y. wrote the manuscript with edits from other authors. All experiments were conducted under the supervision of F.G. All authors read and approved the final manuscript.

