## Supplementary material for "Transdermal diagnosis and therapy using an integrated acoustofluidic patch": SI

**Supporting information:**

**List of Contents**

**Supplementary Figures**

Supplementary Fig. 1 Electronic architecture of the closed-loop system.
Supplementary Fig. 2 Angle-dependent SAW-induced acoustic streaming in the microchannel.
Supplementary Fig. 3 Skin recovery following acoustofluidic patch application.
Supplementary Fig. 4 Heating effect under acoustic actuation.
Supplementary Fig. 5 Acoustofluidic transdermal sampling.
Supplementary Fig. 6 Correlation between histamine levels in serum and interstitial fluid (ISF).
Supplementary Fig. 7 Physiological responses during anaphylaxis in a mouse model.
Supplementary Fig. 8 Respiratory rate dynamics with and without acoustofluidic patch treatment.

**Supplementary Videos**

**Supplementary Movie. 1** Acoustic streaming and nebulization at the outlet of the acoustofluidic injector

**Source data**

The source data file for all the graphics.

### Supplementary Figures

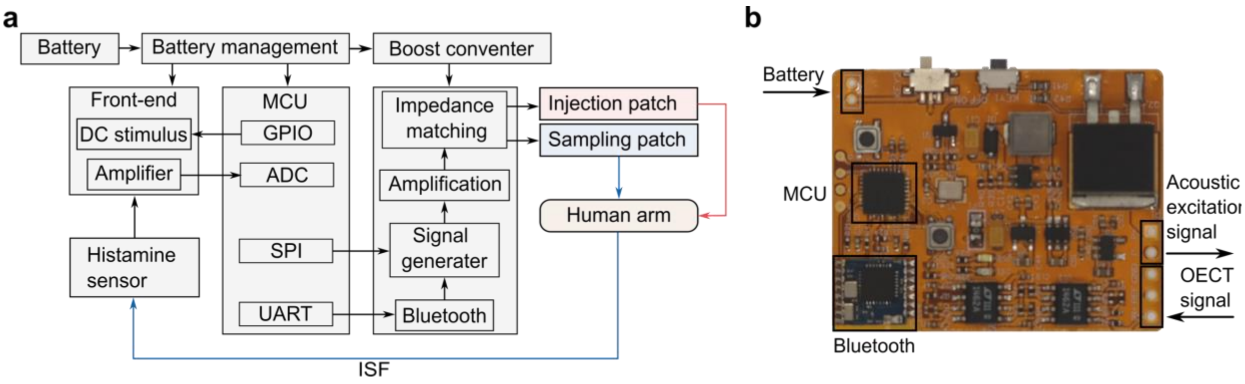


**Supplementary Fig. 1 Electronic architecture of the closed-loop system.** (**a**) Block diagram of the integrated electronic system, including power management, signal generation and amplification, sensor readout, MCU, and wireless communication for coordinated operation of the sampling and injection patches. (**b**) Photograph of the custom-designed printed circuit board (PCB) highlighting key components, including the battery interface, MCU, Bluetooth module, and output ports for acoustic excitation and OECT signal acquisition. MCU, microcontroller unit; ADC, analog-to-digital converter; SPI, serial peripheral interface; UART, universal asynchronous receiver-transmitter; OECT, organic electrochemical transistor; ISF, interstitial fluid.

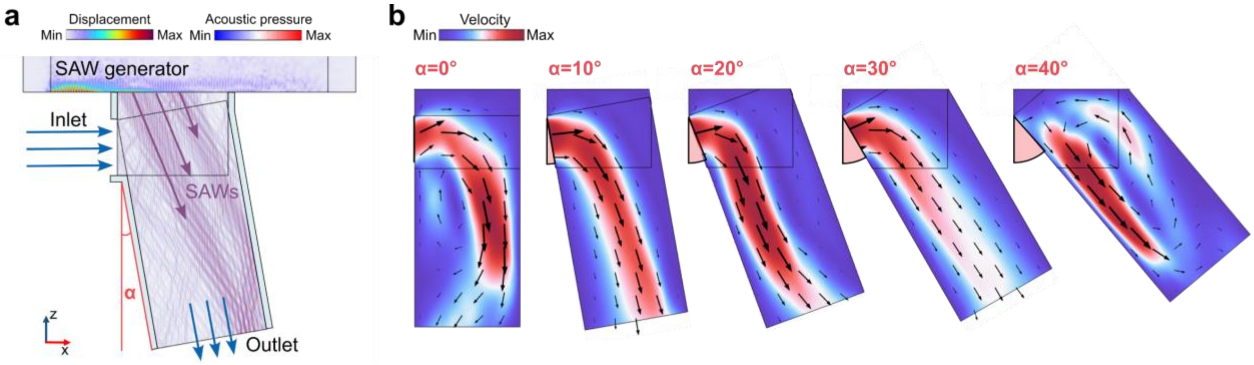


**Supplementary Fig. 2 Angle-dependent SAW-induced acoustic streaming in the microchannel.** (**a**) Finite-element simulation illustrating SAW coupling into the microchannel and the definition of the incident angle (α). (**b**) Simulated velocity fields and streamlines at different SAW incident angles (0°, 10°, 20°, 30°, and 40°). (**c**,**d**) Velocity profiles at the outlet (c) and inlet (d) regions of the microchannel, showing angle-dependent modulation of flow magnitude.

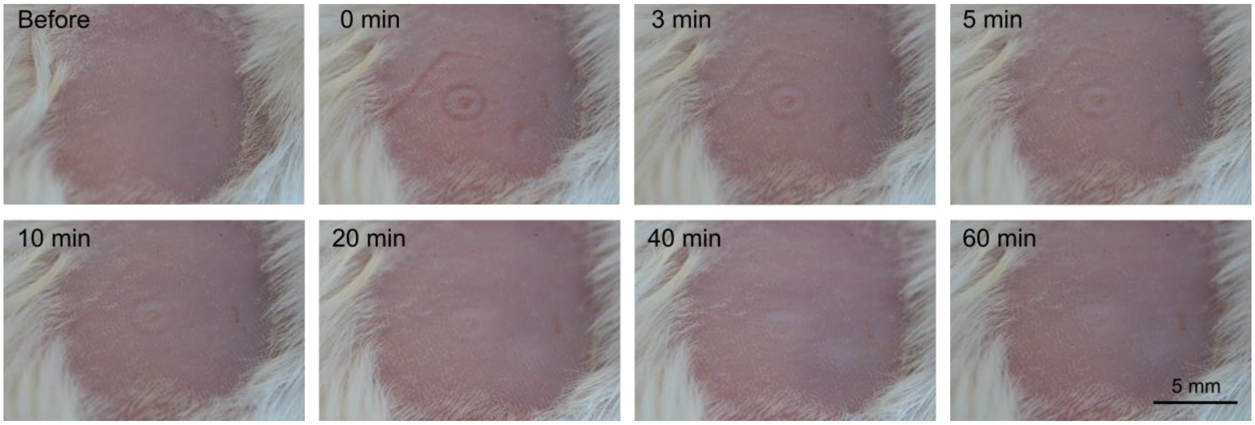


**Supplementary Fig. 3 Skin recovery following patch application**. Time-lapse images of the skin surface before patch application and at 0, 3, 5, 10, 20, 40, and 60 min after patch removal, showing gradual recovery of the skin appearance over time (scale bar, 5 mm).

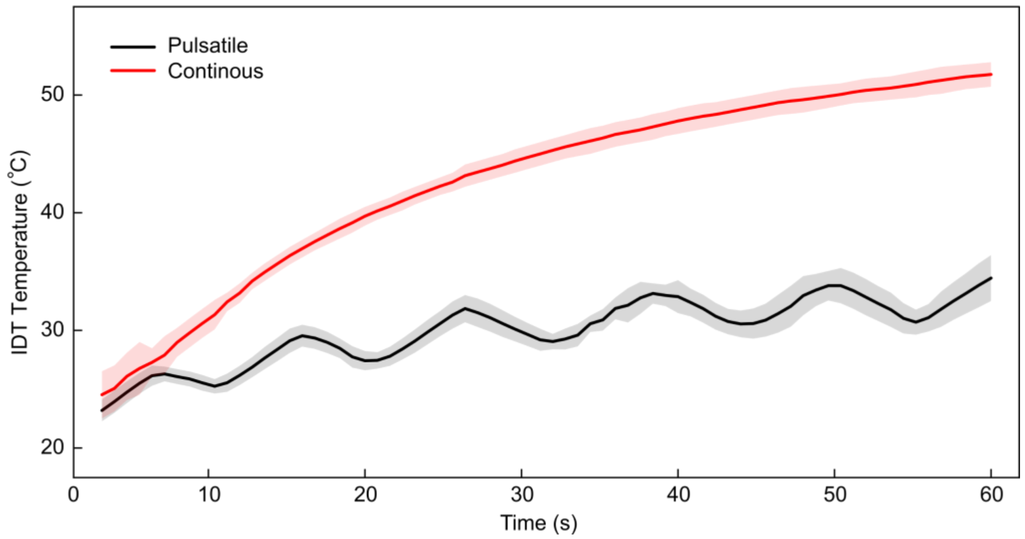


**Supplementary Fig. 4 Heating effect under acoustic actuation.** The temperature of fluid in the chamber over time during operation at 37.4 MHz (Vpp = 1.7 V). The duty cycle of the pulsatile signal is 50%. Data are mean ± s.e.m. (continuous, n=3; pulsatile, n=4).

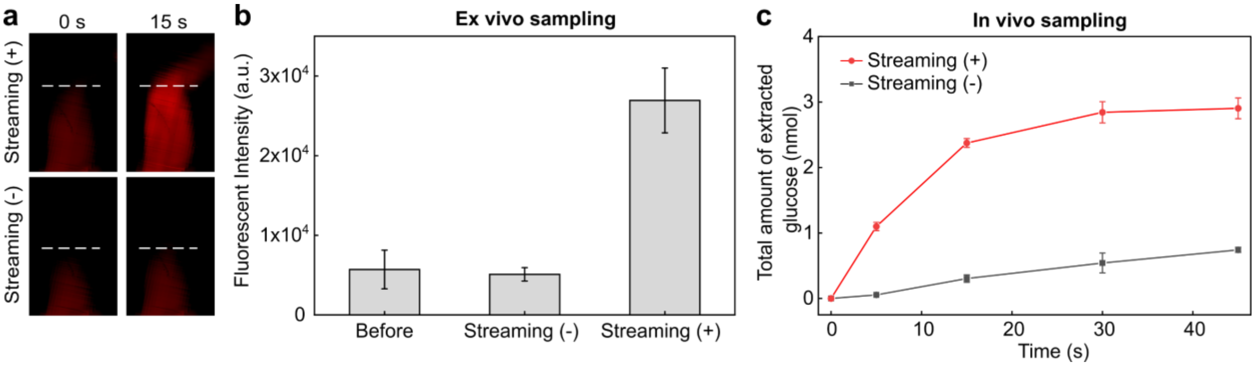


**Supplementary Fig. 5 Acoustofluidic transdermal sampling.** (**a**) Representative fluorescence images showing ex vivo transdermal sampling with streaming OFF and ON at different time points. (**b**) Quantification of fluorescence intensity from ex vivo sampling under different acoustic conditions. Data are presented as mean ± SD; n = 5 independent groups; three positions were measured per group. (**c**) Time-dependent ex vivo sampling performance with and without acoustic actuation, demonstrating enhanced analyte extraction under acoustics ON conditions. Data are presented as mean ± SD; n = 4 independent mice.

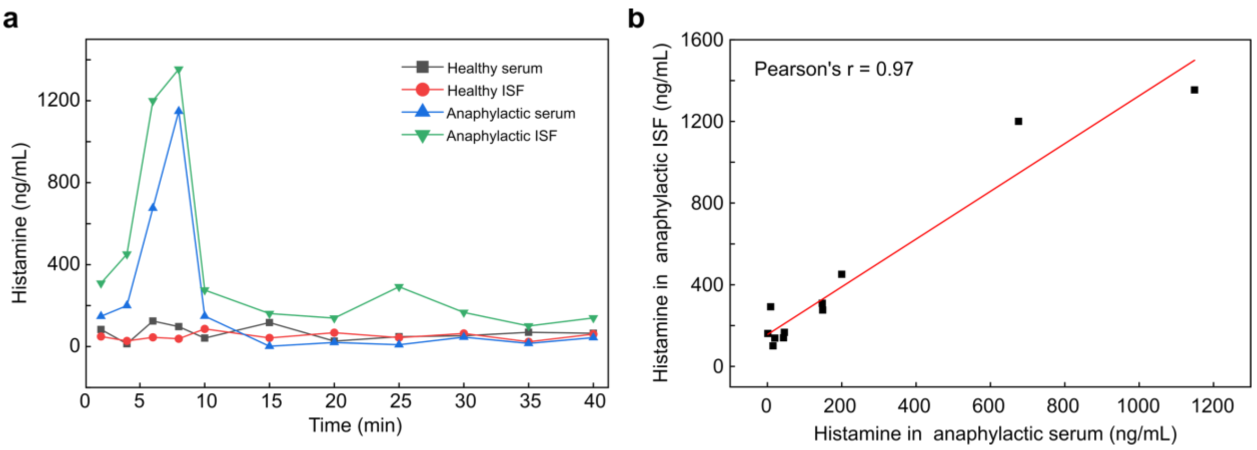


**Supplementary Fig. 6 Correlation between histamine levels in ISF and serum.** (**a**) Time-course profiles of histamine levels measured in serum and interstitial fluid (ISF) under healthy and anaphylactic conditions. (**b**) Correlation analysis between histamine concentrations in anaphylactic ISF and serum, showing a strong positive correlation (Pearson's r = 0.97).

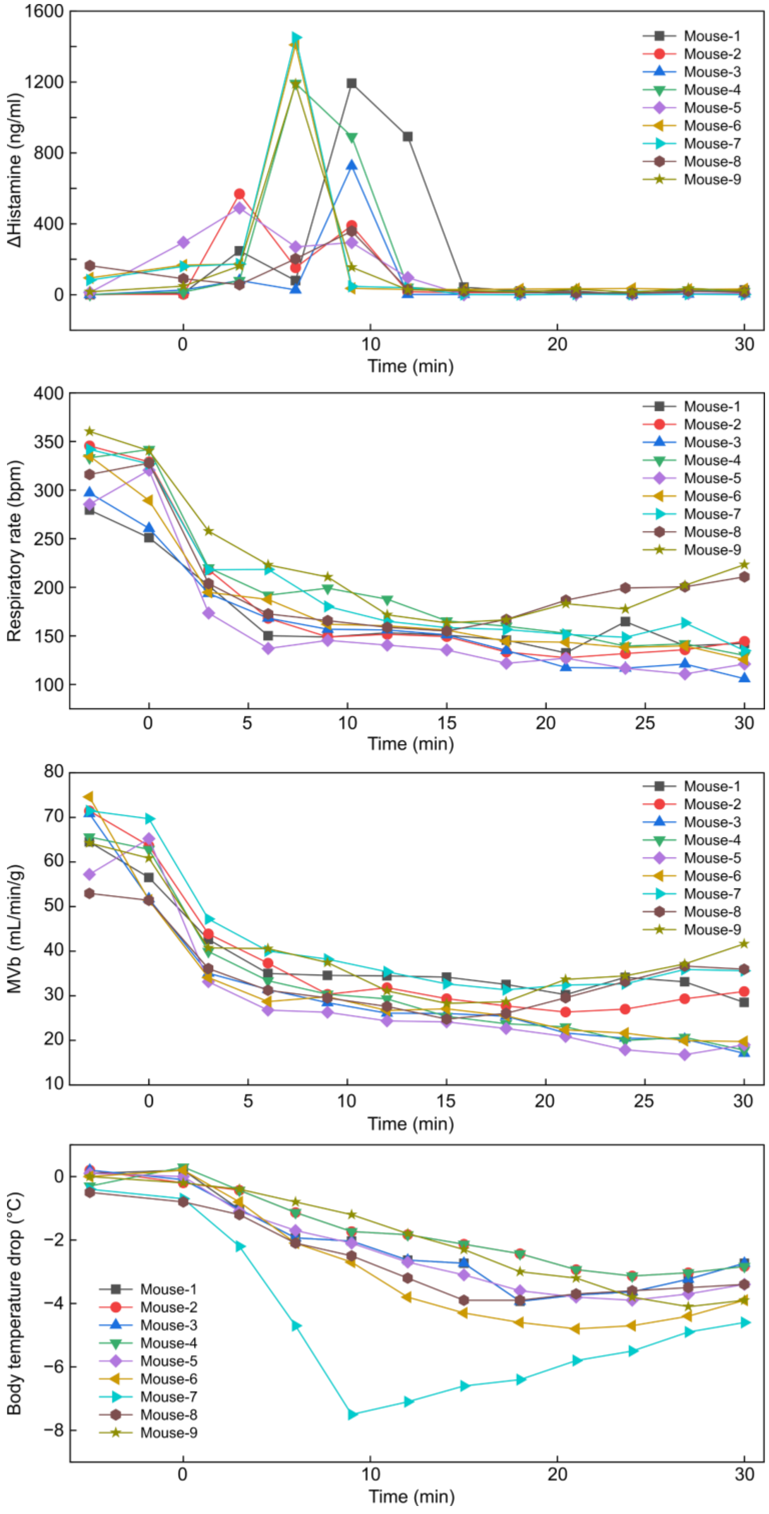


**Supplementary Fig. 7 Physiological responses during anaphylaxis.** Time-course measurements of (**a**) changes in histamine levels, (**b**) respiratory rate, and (**c**) body temperature for individual mice following anaphylactic challenge, illustrating inter-animal variability and the dynamic progression of systemic responses.

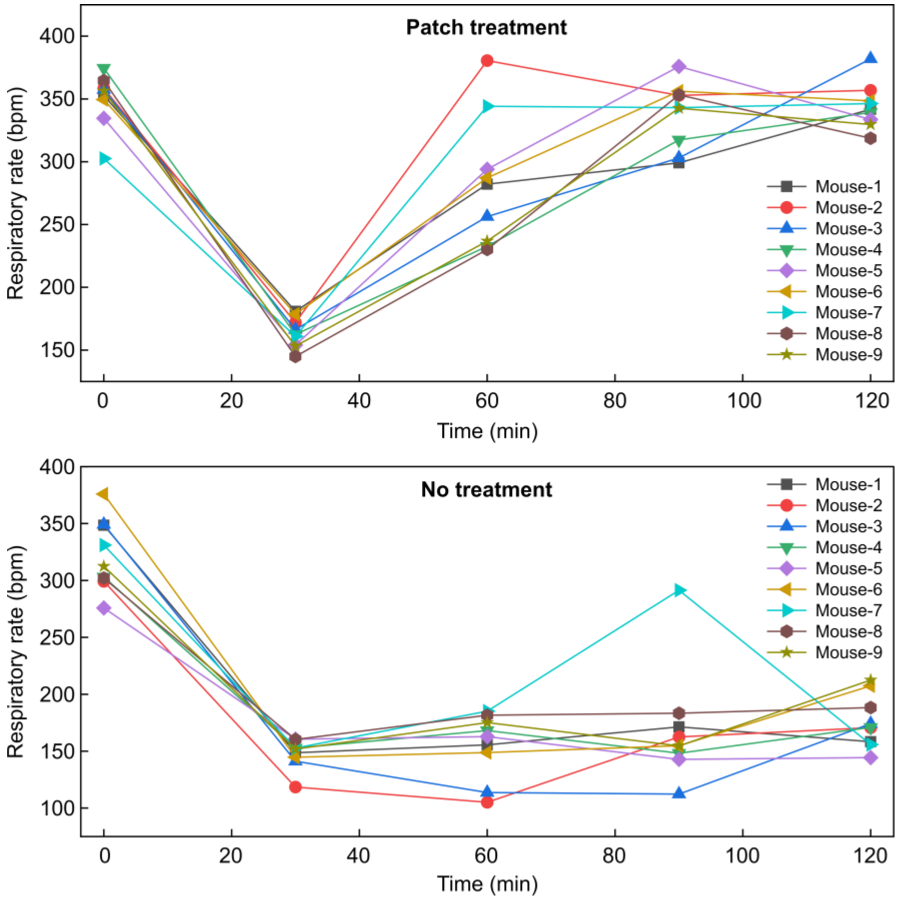


**Supplementary Fig. 8 Respiratory rate dynamics with and without patch treatment.** Time-course measurements of respiratory rate for individual mice subjected to acoustofluidic patch treatment (top) or no treatment (bottom) after anaphylaxis induction, illustrating inter-animal variability in respiratory recovery over time.
